# Using auxotrophic donor strains to explore pQBR57 plasmid host range among environmental soil bacterial isolates

**DOI:** 10.64898/2026.02.11.702040

**Authors:** Alejandro Marquiegui-Alvaro, Anastasia Kottara, Matthew J. N. Thomas, Alberto Scarampi, Micaela Chacón, Michael Brockhurst, Neil Dixon

## Abstract

Plasmid host range (PHR) plays a key role in the spread of ecologically important genes, alongside applications in microbiome engineering, and environmental biotechnology. PHR is a complex trait arising from the combination of plasmid, donor and recipient properties. Most studies of PHR use a single donor strain, leaving the role of the donor unexplored, and often require genetically tagged recipient strains for counter selection, which limits use of non-genetically tractable strains. Here we developed a PHR screening method using auxotrophic donors that bypasses the need to genetically tag recipients, thus allowing the screening of culturable environmental bacterial strains. Specifically, we used two auxotrophic donors (*P. fluorescens* and *P. putida*), and the plasmid pQBR57-tphKAB, an environmental plasmid engineered for terephthalic acid bioremediation. We screened a library of 101 soil isolates, as potential recipients, including common soil genera of soil bacteria, *Pseudomonas, Bacillus* and *Xanthomonas*. We only observed conjugation into other *Pseudomonas*, but donor identity affected PHR, with *P. fluorescens* conjugating the plasmid into more recipient strains than *P. putida*. Phylogenomic analysis revealed that transconjugants clustered with *P. citronellosis* and *P. putida* lineages. In strains that were close relatives of transconjugants but who were unable to acquire the plasmid, we observed 5 defence systems not present in transconjugants that may act as barriers to plasmid acquisition. Our method provides a rapid, tag-free framework for screening PHR in environmental isolates and for investigating the influence of donor identity on plasmid conjugation.

## Introduction

Conjugative plasmids, extrachromosomal genetic elements that encode the genes necessary for their own transfer, are the main driver of horizontal gene transfer in bacterial communities^1^. Conjugation spreads ready-to-use adaptive phenotypes (e. g. xenobiotic metabolism^2^, antibiotic resistance^3^, toxin resistance^4^) across taxonomic boundaries enabling rapid adaptation to changing environmental conditions^5^. Conjugation requires physical contact between donor and recipient cells allowing plasmid transfer via the conjugation pilus^6^. A fundamental limit on plasmid transmission is plasmid host range (PHR), which is predicted to be determined by the combination of microbial host and plasmid genetics^7,8^. PHR is poorly understood for many plasmids^9^. Based upon the taxonomic distribution of hosts, plasmids are often categorised into broad or narrow host range^10^. Broad host range plasmids occur in distantly related taxa^11^, even across multiple kingdoms^12,13^, while narrow host range plasmids are confined within a species or genus^14^. While PHR is encoded by genetic factors of the plasmid itself^15^, it is also affected by the identity of the donor, meaning that the same plasmid in different donors can display differences in the taxonomic identity and range of plasmid recipients^16,17^. Recipients may also affect PHR through mechanisms including plasmid-mediated entry exclusion^18^, defence systems targeting foreign DNA (restriction modification^19^, CRISPR-Cas^20^) or altering the cell surface^21,22^ that prevent plasmid transfer. Given this complex interplay of donor-plasmid-recipient factors upon PHR, it has proven challenging to predict PHR computationally^23^. Nonetheless, understanding PHR is crucial for clinical surveillance of multidrug resistance plasmids and for selecting plasmids to use in biotechnological applications where tailored delivery of genes is required^24–26^. New experimental tools to quantify PHR are needed, particularly for use with non-model bacterial taxa.

Traditional methods for observing plasmid conjugation use donor and recipient strains that have both been tagged with distinct selectable marker genes (e.g., an antibiotic resistance cassette). Transconjugants are then recovered by plating on dual selection media, selecting for both the recipient’s chromosomal marker gene and the plasmid-encoded trait^27,28^. Such methods are of limited value for studying PHR in natural isolates because they require genetic manipulation of recipient strains, which is often challenging outside of a limited number of model organisms^29^. An alternative approach is to counter-select against the donor strain, bypassing the necessity of genetically modifying recipients. Fluorescently-tagging the donor strain and plasmid with distinct fluorophores allows for donors to be separated from transconjugants on the basis of fluorescence using cell sorting^7,30^. This method is powerful because it can be used with unculturable organisms but requires access to expensive state-of-the-art equipment (e.g. FACS). A lower-tech alternative means of counter-selection is to use an auxotrophic donor (i.e. a strain unable to grow in the absence of an essential nutrient). Here, donors can be separated from transconjugants by plating on media that lacks the relevant nutrient but is supplemented with a selective agent for the plasmid-encoded trait. For example, *E. coli* strains with diaminopimelic acid (DAP) auxotrophy have been used to transfer cloning plasmids (derivatives of pECE743) into *Bacillus*^31^ and the integrative pSET152 plasmid to *Actinomycetes* strains^32^. In addition, a D-alanine *B. subtilis* auxotroph was used to transfer an integrative and conjugative element to a range of Gram positive bacteria^33^.

In this study, we develop a high-throughput PHR screening method using *P. fluorescens* SBW25 Δ*panB tdTomato* (pantothenate auxotroph) and *P. putida* KT2440 Δ*trpD gfp* (tryptophan auxotroph) donors along with the environmental plasmid pQBR57-KAB (**Fig 1**). pQBR57 is a conjugative megaplasmid (307kb) isolated from the sugar beet phytosphere^34^ that naturally encodes a mercury resistance operon within the Tn5042 transposon^28^. The transfer, maintenance and fitness cost properties of pQBR57 within *P. fluorescens* have been thoroughly characterized^35–38^. pQBR57-tphKAB (hereafter pQ-KAB) is an engineered version of pQBR57 equipped with transport and catabolic genes for the assimilation of terephthalic acid (tphKAB)^39^. Terephthalic acid is the aromatic monomer of polyethylene terephthalate (PET) plastic. pQ-KAB has shown promise as a potential vector for genetic bioaugmentation-mediated bioremediation of soils^40^. We assessed pQ-KAB’s PHR from each donor into a panel of recipients comprising 101 diverse bacterial isolates isolated from potting soil. We show that our method can be used to rapidly screen for plasmid transfer to unlabelled culturable environmental soil bacterial. Moreover, we show that PHR varies between donor strains and identify variation in defence systems among potential recipients that may limit plasmid acquisition.

**Figure 1.**
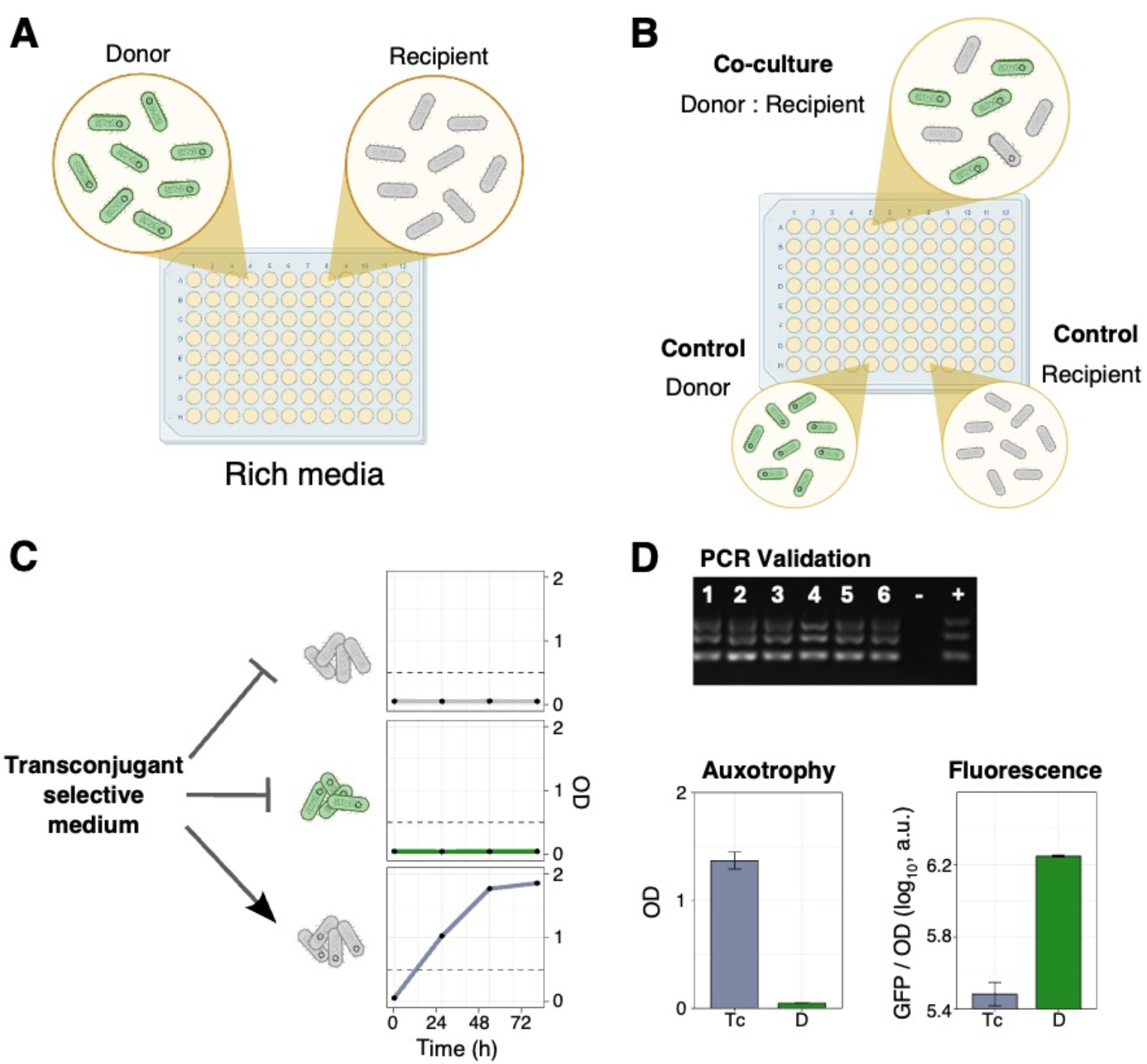
Method workflow. A representative example using *P. putida* KT2440 Δ*trpD gfp* as the donor of pQ-KAB and *Pseudomonas* isolate D5 as the recipient. (**A**) Donor and recipient are grown independently in rich media, then (**B**) they are mixed in co-culture in rich media, maintaining monocultures as controls. (**C**) After 24-hour incubation, co-culture are re-inoculated at three dilutions in transconjugant-selective media (Tc-media, M9 10 mM glucose 20μM HgCl_2_), the lack of essential nutrient inhibits donor growth while mercury prevents plasmid-free cells from growing, thus selecting for transconjugants. Wells with optical density (OD) above 0.4 are streaked on plasmid-selective media (KB agar 20μM HgCl_2_). (**D**) Transconjugants confirmed by plasmid presence by PCR screening, growth on M9 10 mM glucose (no Trp supplementation) and lack of fluorescence.

## Results

### Engineering and validation of auxotrophic donor cells

We first characterised the auxotrophic strains we used as donors. The *P. fluorescens* pantothenate auxotroph, *P. fluorescens ΔpanB* (hereafter PF Δ*panB*), was originally constructed by homologous recombination^41^. Production of panthothenate (vitamin B5) begins with PanB, a ketopantoate hydroxymethyltransferase which catalyses the conversion of α-ketoisovalerate to α-ketopantoate. We validated the pantothenate auxotrophy by quantifying growth in minimal medium supplemented with a range of pantothenate concentrations (**Fig 2A**), confirming that PF Δ*panB* is unable to grow at concentrations below 20 nM. We constructed the *P. putida* tryptophan auxotroph, *P. putida ΔtrpD* (hereafter PP Δ*trpD*), by homologous recombination with a suicide vector^42^. Tryptophan is synthesised from chorismate via the action of *trpDC* operon^43^, where TrpD encodes an anthranilate phosphoribosyltransferase which catalyzes the essential step in tryptophan biosynthesis^44^. We validated the auxotrophy by quantifying growth in minimal medium supplemented with a range of tryptophan concentrations (**Fig 2B**), confirming that PP Δ*trpD* is unable to grow at concentrations below 1μM. Both auxotrophic donors, PF Δ*panB* and PP Δ*trpD*, were then chromosomally tagged with tdTomato or GFP, respectively, under a constitutive promoter. pQ-KAB was then conjugated into each donor from *P. fluorescens* SBW25.

**Figure 2.**
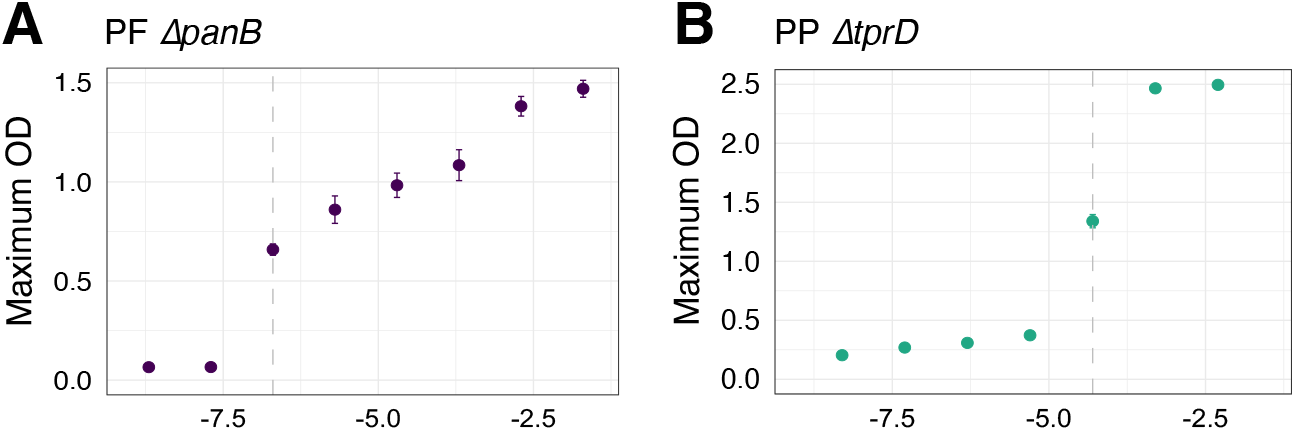
Auxotroph confirmation. (**A**) Pantothenic acid auxotrophy of *P. fluorescens ΔpanB* (PF Δ*panB)* in a gradient of pantothenic acid concentrations, (**B**) Tryptophan auxotrophy of *P. putida ΔtrpD* (PP Δ*trpD*) in a gradient of tryptophan concentrations; both in minimal media supplemented with 10 mM glucose for 24 hours at 30°C.

### Taxonomic diversity of soil isolates used as potential recipient panel

We isolated 128 pure colonies from King’s B (KB) agar plates inoculated with potting soil wash. Prior work suggests that this method captures a diversity of culturable bacterial strains^45^. The isolates were then screened for ability to grow in minimal media with glucose and sensitivity to Hg^2+^ to filter out isolates unable to grow in the culture conditions of our screen or with native mercury resistance. These requirements ensured that only plasmid-carrying recipients could grow in transconjugant-selective media (Tc-media, M9 10 mM glucose 20μM HgCl_2_) due to recipients’ sensitivity to mercury and the donors’ auxotrophies (**Fig 3**). These requirements resulted in 101 isolates that were taxonomically characterized by sequencing the full 16S rRNA gene (**Fig 3**). 64 isolates belonged to *Pseudomonas*, while 37 belonged to a diverse range of bacterial taxa including *Bacillus, Pandorea, Rhodanobacter, Xanthomonas* or *Brucella*. These potential recipient strains are representative of the culturable fraction of soil bacterial diversity^46,47^.

**Figure 3.**
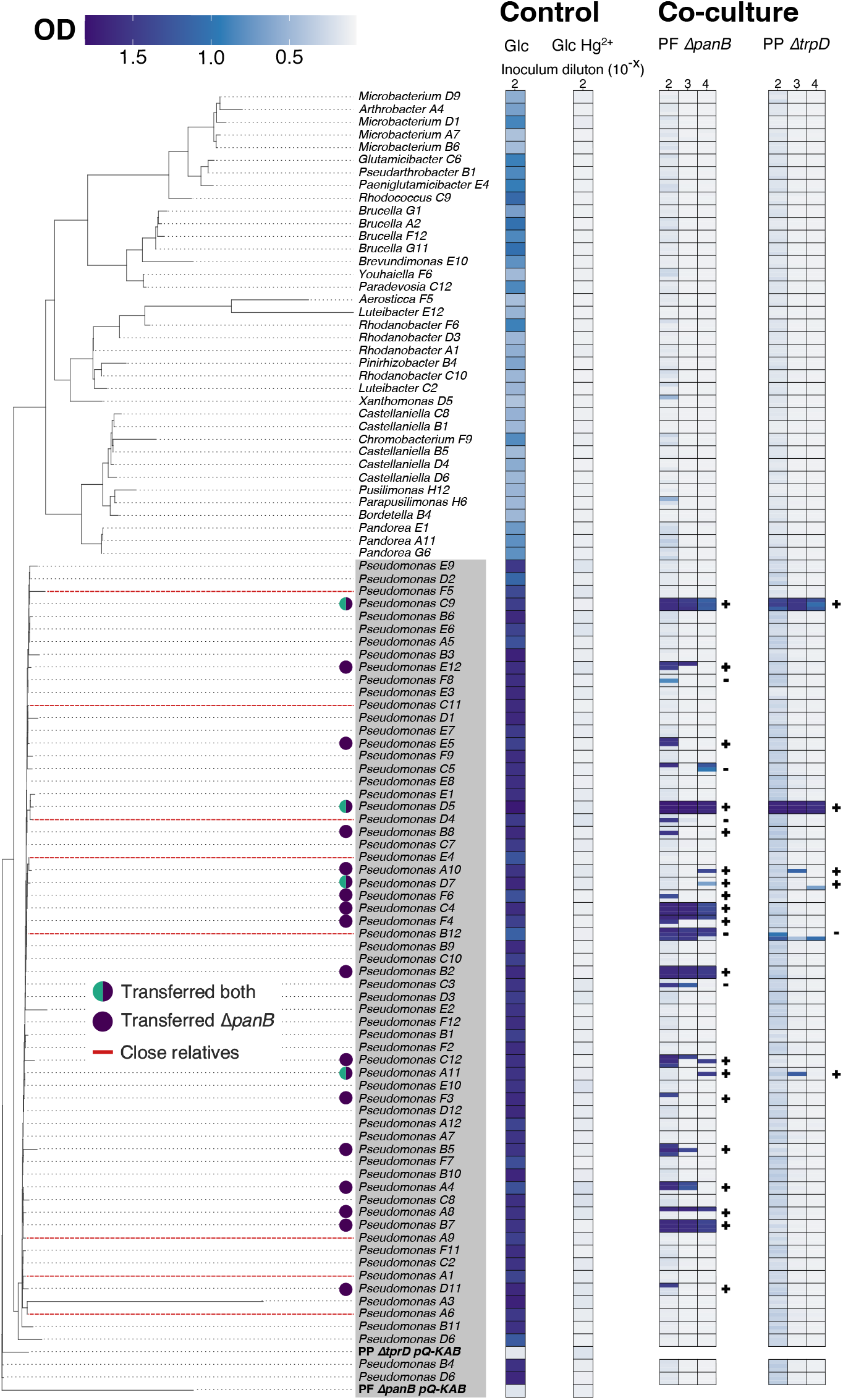
Phylogenetic tree of isolates and growth in transconjugant-selective media. Phylogenetic tree of soil isolates based on full 16s rRNA sequence, branch length has been scaled down for non-*Pseudomonas* and PF Δ*panB* to help visualization. The grey area represent *Pseudomonas* genus. Purple and bicolour circles show the detected transconjugants from PF Δ*panB* or both donors respectively, close relatives that did not acquire the plasmid are marked with red dotted line. The heatmaps adjacent to the tree represent the maximum OD in a 72-hour incubation. The first heat map column is from monocultures of each isolate and donor in M9 10 mM glucose (positive control), followed by monocultures in transconjugant-selective medium M9 10 mM 20 μM HgCl_2_ (negative control). Then, the three columns under PF Δ*panB* represent the co-culture of isolate and corresponding donor, at three inoculation dilutions in transconjugant-selective media, while the remaining three are the coculture of each isolate and PP Δ*trpD* at three inoculation dilutions. For the coculture heatmaps, three replicates are shown for each isolate-donor combination. Next to the putative transconjugant growth, the outcome of the PCR test is displayed.

### pQ-KAB preferentially conjugates into Pseudomonas recipients

Our PHR screen identified 6 putative transconjugants from PP Δ*trpD* and 23 from PF Δ*panB* (i.e., growth exceeded threshold OD_600_ > 0.4) (**Fig 3**, co-culture heat maps). All of the transconjugants belonged to *Pseudomonas*. PCR validation of putative transconjugant colonies suggested that several had not actually gained the plasmid. In total, we were able to confirm 4 transconjugants from PP Δ*trpD* and 19 transconjugants from PF Δ*panB*. Thus, PF Δ*panB* transferred the plasmid to 29.7% of the *Pseudomonas* isolates (19/64) and PP Δ*trpD* to 6.3% (4/64) (**Fig 4A** and **4B**). The PHR of pQ-KAB from PP Δ*trpD* was a subset of the PHR observed from PF Δ*panB* (*Pseudomonas*-A11, *Pseudomonas*-C9, *Pseudomonas*-D5 and *Pseudomonas*-D7, marked with a bicolour circle in **Fig 3**). Plasmid transfer success did not vary with phylogenetic distance of the recipient from the donor (logistic regression, p-value = 0.17 for PF Δ*panB* and p-value = 0.26 for PP Δ*trpD*).

**Figure 4.**
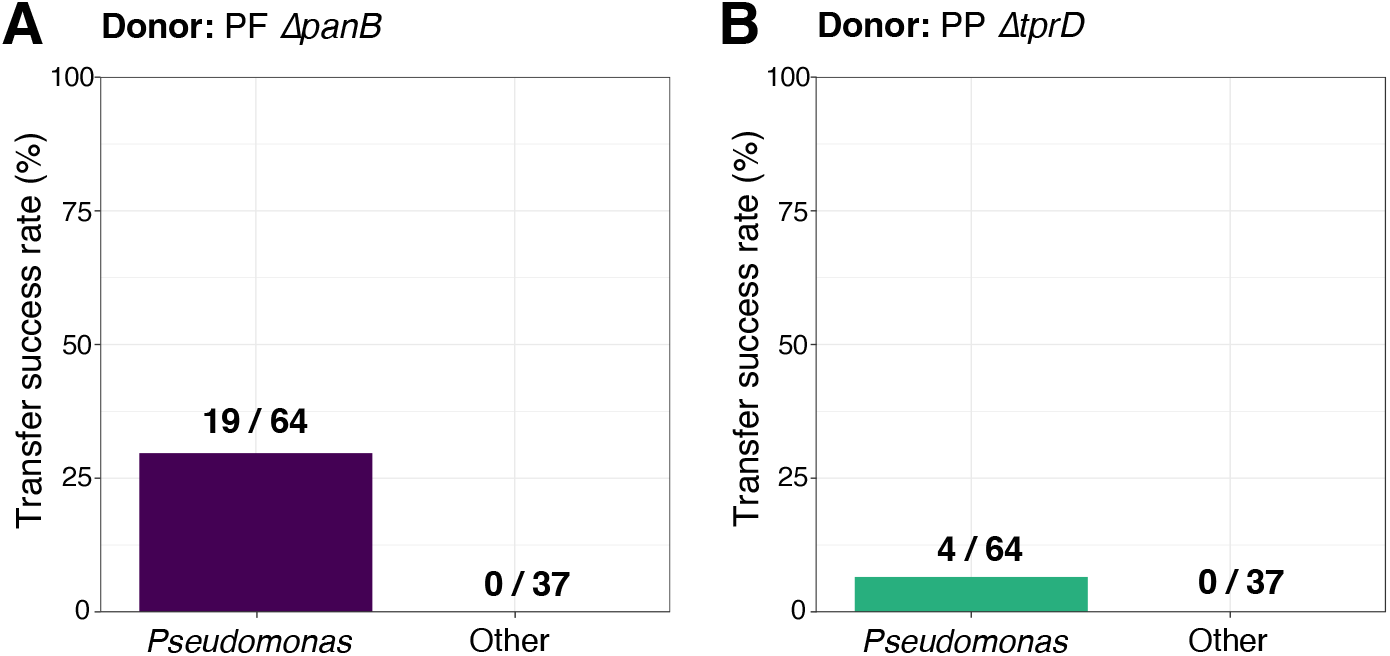
Transfer success rate is donor-dependent. **(A)** Transfer success rate within-genus (*Pseudomonas*) and outside-genus (non-*Pseudomonas*) with PF Δ*panB* pQ-KAB as the donor and (**B**) with PP Δ*trpD* pQ-KAB.

### Transconjugants lack specific defence systems compared to close relatives

To identify transconjugants to species-level, we obtained whole genome sequences for each of the 19 PCR-confirmed transconjugants. We also obtained whole genome sequences for 8 isolates that were phylogenetically closely related to transconjugants (according to their 16s rRNA sequence) but were unable to acquire pQ-KAB from either donor in our PHR screen (**Fig 3**). Combining these genomes with representative genomes from each of the 14 Pseudomonas clades^48^ we constructed a core-genome phylogeny using 71 single-copy genes (**Fig 5A**). Eighteen of the 19 transconjugants grouped with *Pseudomonas citronellolis*, a bacterium found in soil and on plant surfaces^49^, with the remaining transconjugant, *Pseudomonas*-D5, clustering with *P. putida*. We compared the defence system content of transconjugants with close relatives that could not acquire the plasmid. Although the groups did not differ in the number of defence systems (paired sample t-test, p-value = 0.33) (**Fig 5B**), five defence systems were found exclusively in the close relatives that could not acquire the plasmid (**Fig 5C**). Specifically, these were the characterised anti-phage defence systems Erebus, Nantosuelta and Zorya, as well as two defence systems with unknown mechanisms (Gao_Upx and DS-15).

**Figure 5.**
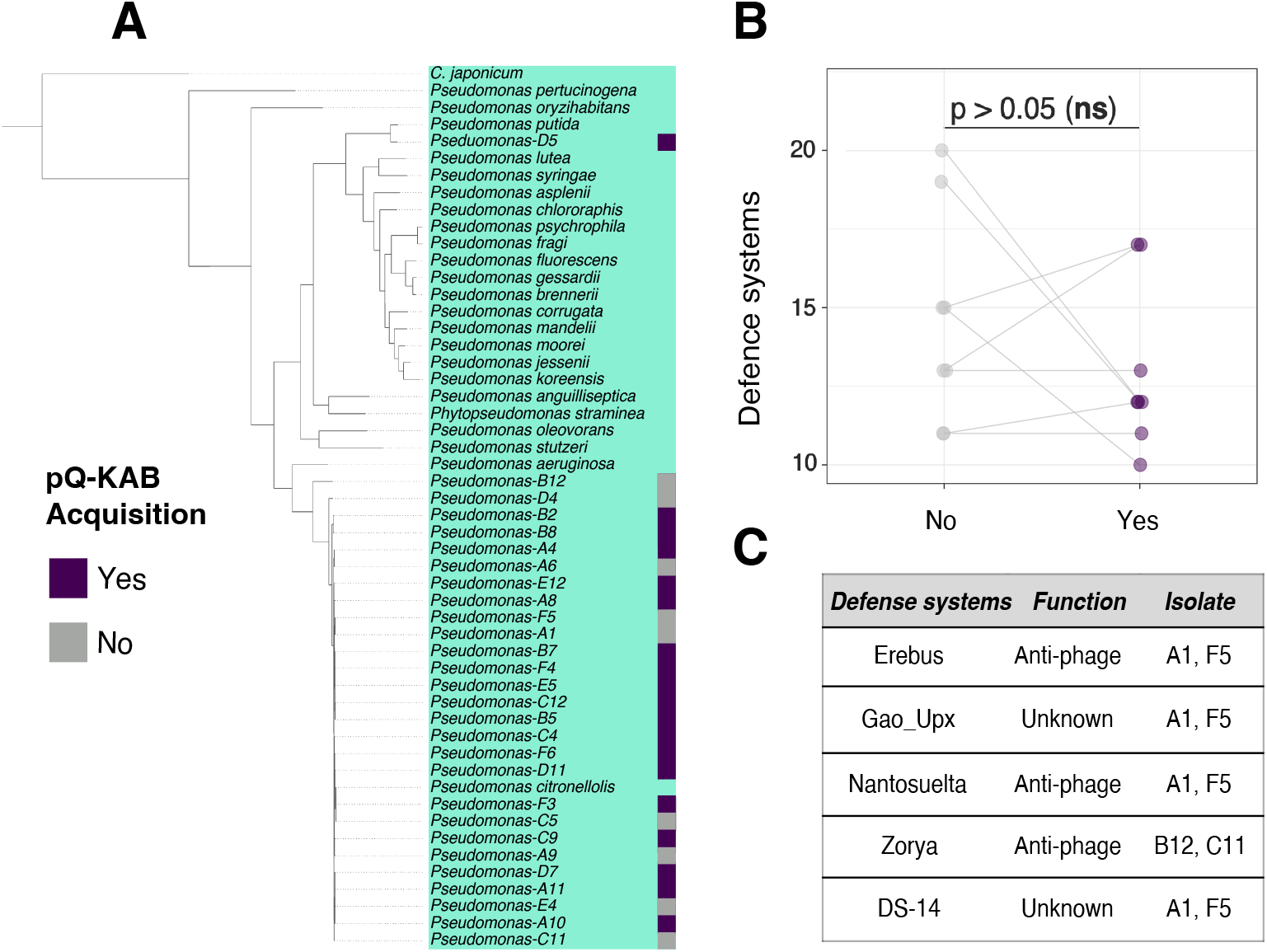
Taxonomic identification of transconjugants and closely related neighbours, and their defence systems. (**A**) Phylogenomic tree of pQ-KAB transconjugants, closely related isolates that did not acquire the plasmid, and representative genomes from each *Pseudomonas* clade with the tree rooted with *C. japonicum* as an outgroup, similar to Gomilla et al (2015)^48^. (**B**) Total number of defence systems identified with *defense-finder*^50,51^ between the isolates that could acquire the plasmid and their close relatives, significance from paired sample t-test. (**C**) Defense systems identified exclusively in the isolates that could not acquire the plasmid, and their putative function.

### Discussion

Here we report the development of a PHR screening method using two auxotrophic donors that enabled screening of 101 soil-isolated environmental bacteria for their ability to acquire pQ-KAB by conjugation. We identified 19 strains of *Pseudomonas* capable of acquiring pQ-KAB, without the need to genomically tag any of the recipient strains. Our data shows that pQ-KAB’s PHR is likely limited to within the *Pseudomonas* genus. In addition, pQ-KAB’s PHR was affected by the donor identity (**Fig 4A** and **B**) and the defence systems encoded by potential recipients, which may limit pQ-KAB transfer.

Our findings corroborate previous experiments using wild-type pQBR57 showing that these plasmids preferentially conjugate into *Pseudomonas* species^36,52^. In contrast, a study by Hall et al (2020)^35^ using *P. fluorescens* as the donor of pQBR57 reported, via epicPCR targeting the *merA* gene, the transfer of the mercury-resistance cassette beyond *Pseudomonas*, including Burkholderiales, Rhizobiales, and even Bacillales. This broader host range could be due to the target gene, *merA*, being located on a transposon (Tn5042). This could have enabled *merA* to relocated to the chromosome during transient plasmid acquisition by these species or onto other mobile genetic elements that these taxonomic orders can then acquire^52,53^. In addition, our findings emphasize the previously reported role of donor identity upon PHR^17^. *P. putida* has a lower stability and conjugation rate of pQBR57 when compared to *P. fluorescens*^54^, which could explain the difference in their ability to transfer pQ-KAB (**Fig 4**). Our data suggest that *P. fluorescens* is a more efficient plasmid donor, consistent with its role as a plasmid reservoir in co-culture with *P. putida* in potting soil^37^.

In general, the likelihood of conjugation is expected to decline with increasing phylogenetic distance between donor and recipient^55^. Although we observed that pQ-KAB appears restricted to *Pseudomonas*, we observed no significant phylogenetic signal in PHR. Such patterns are likely to be more apparent at higher levels of taxonomic classification (e.g. class, orders)^55^. At lower taxonomic levels (e.g., genus), phylogenetic distance shows a less strong correlation with the likelihood of transfer^56^. Within genus, the presence of defence systems^56^ or other plasmids^18^ appear to be more important determinants for the success of plasmid transfer. Within *Pseudomonas*, most of the pQ-KAB transconjugants clustered within the species *P. citronellolis* (**Fig 5A**), suggesting that pQ-KAB is a relatively narrow host range plasmid. Moreover, the findings suggest that the natural host of the parental pQBR57, which is unknown due to its isolation by exogenous capture from a sugar beet field^34^, may have been a strain of *P. citronellolis*.

Comparison of defence systems in closely related strains that did or did not receive pQ-KAB supported a potential role for bacterial immunity in PHR here. Although the total number of defence systems did not differ between transconjugants and non-transconjugants (**Fig 5B**), we identified five systems that were exclusively found in close relatives unable to acquire pQ-KAB (**Fig 5C**). This dissociation could indicate targeting of pQ-KAB by these defence systems. Where known, the mechanism of action of these defence systems (e.g. Zorya) is to digest foreign nucleic acid via nuclease effectors^57–59^. Although some defence systems are thought to exclusively target phage DNA and have no effect on conjugation (e.g. Zorya^59^), there are other systems where the specificity of targeted MGEs remains unclear. That defence systems impose selection on conjugative plasmids is clear from plasmids encoding anti-defence systems in their leading region^60^. Our data potentially suggest that defence system associated nucleases could interfere with the leading strand of pQ-KAB upon conjugation, thus disrupting the conjugation process.

Our PHR screening method has several advantages relative to existing conjugation protocols. First, it bypasses the need to genetically manipulate the recipient, enabling testing of plasmid host range in environmental bacteria. As such, this protocol could be used to identify plasmid-donor pairs capable of transferring genetic material to non-model organisms for investigating their biotechnology potential^61–63^. Second, our method is relatively high throughput, since it can be carried out in 96-well plates or 364-well plates, with the complete workflow takes only 5 days to perform (**Fig 1**). Third, it is cost-effective and reliant on common lab consumables and equipment (optical density plate reader, incubator, thermocycler). The more expensive techniques used (e.g. whole-genome sequencing) are not essential, although they do provide an additional layer of characterization of transconjugants. The main limitation of our method is that recipient strains must fulfil certain criteria: (i) lab cultivability; (ii) sensitivity to the plasmid selective agent; (iii) prototrophic growth. In addition, we observed a low rate of false positives whereby non-plasmid carrying recipients grew in transconjugant-selective media. We hypothesize that growth in these co-cultures may be the result of disruption of cell membrane integrity by mercuric ions^64^, which could release sufficient pantothenate or tryptophan to allow donor growth. However, PCR validation for the plasmid can rapidly filter out such false positives.

Overall, our PHR screening method provides a robust and accessible platform for assessing the effect of donor on PHR for panels of non-model environmental isolates without requiring genetic manipulation of potential recipients. By enabling enrichment and isolation of transconjugants, it facilitates downstream genomic and phenotypic characterization of newly formed plasmid–host associations As such, this approach offers a valuable tool for dissecting the ecological and evolutionary constraints shaping plasmid spread, and for identifying donor–recipient and plasmid-recipient combinations with potential utility in environmental microbial biotechnology.

## Materials and Methods

### Donor strain construction

*Pseudomonas fluorescens ΔpanB* SBW25 is a pantothenic acid (vitamin B5) auxotroph obtained is from Rainey et al (1998)^41^. *Pseudomonas putida ΔtrpD* was constructed by using the pK18mobsacB^42^ vector which incorporated a gentamicin cassette at the locus of *trpD* via homologous recombination and selection of double crossover with sucrose counter-selection. *Pseudomonas putida ΔtrpD* was fluorescently tagged with a constitutively expressed GFP cassette using the same delivery vector and integrating at the *fpvA* locus. *Pseudomonas fluorescens* was tagged with a tdTomato-expressing cassette using the mini-Tn7 system^65^ to insert at the chromosomal attTn7 site. The plasmid pQBR57-tphKAB (pQ-KAB) is from Marquiegui-Alvaro et al (2025)^40^ and was horizontally transferred to the auxotroph via conjugation from a non-fluorescent *P. putida* donor. Briefly, after coculturing in King’s B media for 24 hours at 30°C, plated on plasmid-selective media (King’s B agar with HgCl_2_ 20 μM), the colonies with fluorescence were picked and their auxotrophy confirmed by their lack of growth M9 minimal media with 10 mM glucose.

### Microbial soil isolates

Briefly, we added 10 g John Innes No. 2 compost soil in 30 ml universal vials and added 10 ml of M9 salt solution and 20 glass beads. After vortexing for 1 min, the soil wash was plated onto KB agar plates. These plates were then incubated at 30°C with 80% humidity for 48 hours. Subsequently, we selected individual bacterial colonies that we grew in KB media and stored in 25% glycerol at -80°C. We systematically filtered out the bacterial isolates with native resistance to mercury by growing them for 48 hours at 30°C in 96-well microplates with King’s B and 20 μM HgCl_2_.

### Plasmid host range screening

Three biological replicates of each donor (*P. fluorescens ΔpanB tdTomato* pQ-KAB and *P. putida ΔtrpD gfp* pQ-KAB) were picked by streaking in King’s B agar with 20 μM Hg (II). These were grown in 5ml of King’s B with 20 μM Hg (II) at 30^°^C for 18 hours at 180 rpm. The cultures were washed with King’s B to remove any traces of Hg (II). For the soil isolates, they were grown in King’s B in 96-round well plates at 30^°^C for 18 hours. Donors and recipients were inoculated at a 1:5 colony forming unit ratio in favour of recipients in King’s B and grown at 30^°^C for 18 hours. Then, the plates were washed with minimal media to remove any traces of essential nutrients (pantothenic acid and tryptophan) and inoculated in selection media - minimal media with 10 mM Glucose and 20 μM Hg (II) – at four dilutions (neat, 1/10, 1/100 and 1/1,000). These cultures were grown at 30^°^C for 72 hours. The co-cultures that significantly grew, compared to the single-strain controls (i.e. OD > 0.4) were streaked in KB agar ± 20 μM Hg (II) and single colonies were screened for the presence of the plasmid via multiplex PCR targeting the *tphK, uvrD* and *merA* genes of the plasmid. To confirm the absence of donors, we grew them in minimal media with 10 mM Glucose and in KB to re-validate lack of fluorescence.

### Whole genome sequencing and phylogenomic analysis of transconjugants

After confirmation of transconjugants via PCR, to validate plasmid carriage, and lack of auxotrophy and fluorescens, to ensure it was not the donor, single colonies were grown in King’s B media with 20 μM Hg (II) at 30°C for 16 hours and 180 rpm. Cell count was estimated via colony forming units and 5 x 10^9^ cells were washed with M9 media and resuspended in Zymo DNA/RNA Shield. Long-read sequencing with Oxford Nanopore was outsourced to Plasmidsaurus Ltd. The following packages were used in the workflow: flye^66^ (genome assembly), prokka (annotation), anvi’o^67,68^ (phylogenomic of 71 single-copy core genes), FastTree^69^ and iqtree^70^ (tree plotting). For the phylogenomic tree, we reference genomes were downloaded from *Pseudomonas* database (www.pseudomonas.com).

## Supplementary Figures

**Supplementary Figure 1.**
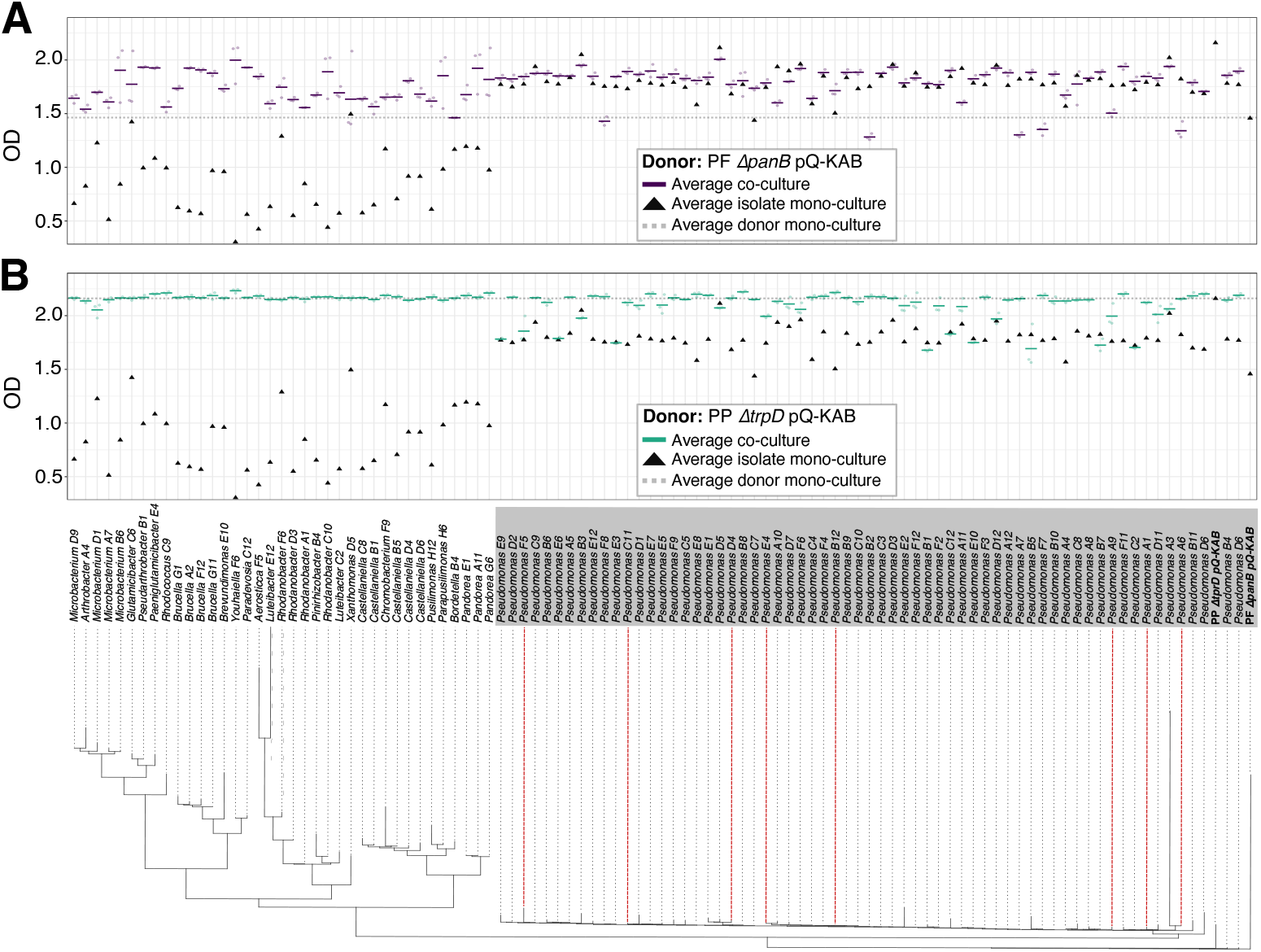
Growth of isolates in mono-culture and co-culture with each donor in KB media. OD reading after 24-hour incubation at 30°C. (**A**) Isolates in co-culture with Δ*panB* in triplicates (purple circles) with the average shown with a horizontal purple line, the dotted line represents the OD of Δ*panB* in mono-culture while the triangles are the OD of each isolate in mono-culture. (**B**) Analogous to (A) but with Δ*trpD* as donor.

## Notes

### Competing Interest Statement

The authors have declared no competing interest.

